# Muscle strength, size and composition following 12 months of gender-affirming treatment in transgender individuals: retained advantage for the transwomen

**DOI:** 10.1101/782557

**Authors:** A Wiik, TR Lundberg, E Rullman, DP Andersson, M Holmberg, M Mandić, TB Brismar, O Dahlqvist Leinhard, S Chanpen, J Flanagan, S Arver, T Gustafsson

## Abstract

**Objectives:** This study explored the effects of gender-affirming treatment, which includes inhibition of endogenous sex hormones and replacement with cross-sex hormones, on muscle function, size and composition in 11 transwomen (TW) and 12 transmen (TM).

**Methods:** Isokinetic knee extensor and flexor muscle strength was assessed at baseline (T00), 4 weeks after gonadal suppression of endogenous hormones but before hormone replacement (T0), and 3 (T3) and 11 (T12) months after hormone replacement. In addition, at T00 and T12, we assessed lower-limb muscle volume using MRI, and cross-sectional area (CSA) and radiological density using CT.

**Results:** Thigh muscle volume increased (15%) in TM, which was paralleled by increased quadriceps CSA (15%) and radiological density (6%). In TW, the corresponding parameters decreased by −5% (muscle volume) and −4% (CSA), while density remained unaltered. The TM increased strength over the assessment period, while the TW generally maintained or slightly increased in strength. Baseline muscle volume correlated highly with strength (R>0.75), yet the relative change in muscle volume and strength correlated only moderately (R=0.65 in TW and R=0.32 in TM). The absolute levels of muscle volume and knee extension strength after the intervention still favored the TW.

**Conclusion:** Cross-sex hormone treatment markedly affects muscle strength, size and composition in transgender individuals. Despite the robust increases in muscle mass and strength in TM, the TW were still stronger and had more muscle mass following 12 months of treatment. These findings add new knowledge that could be relevant when evaluating transwomen’s eligibility to compete in the women’s category of athletic competitions.

## Introduction

Competitive sports have been divided primarily by the concepts of male/female identity. However, these traditional athletic categories do not account for transgender persons who experience incongruence between the gender assigned at birth and their experienced gender identity. The question of when it is fair to permit transgender persons to compete in sport in line with their experienced gender identity is a delicate issue given the desire to ensure fair, safe and meaningful competition while at the same time protecting transgender individuals’ rights and autonomy [1–4]. This has been a highly controversial topic, not least after the recent International Association of Athletics Federations (IAAF) regulations on testosterone limits, which was brought about by the rare genetic condition “differences of sex development” (DSD) [5, 6].

The International Olympic Committee stated in 2016 that Transmen (TM, previously termed female-to-male) are allowed to compete in the male category without restrictions, while Transwomen (TW, previously termed male-to-female) must have a declared gender as female and have testosterone levels below 10 nmol/L for at least 12 months prior to competition [7]. The IAAF in turn states that their expert medical panel shall make a comprehensive review of the athlete’s case and that the athlete is eligible to compete in women’s competition if the panel determines that her medical treatment following sex reassignment has been administered in a verifiable manner and for a sufficient length of time to minimize any advantage in women’s competition [8]. The obvious problem is that there is essentially no data on how long it takes to negate the athletic advantages of many years of male testosterone levels, as TW have experienced prior to commencing gender-affirming treatment. To add further complexity to the issue, the athletic advantage of both endogenous and exogenous testosterone levels is highly debated [9–13].

A meta-analysis reported that TM on average gain 3.9 kg of lean body mass whereas TW lose 2.4 kg during the course of 12 months of cross-sex hormone therapy [14]. Evidence of lower-limb muscle size changes has been provided by a few studies of the first 12 months of treatment in transgender individuals undergoing hormone therapy. A substantial increase in muscle mass (10-19%) with testosterone administration was reported in TM [15–17], while TW undergoing testosterone suppression and estrogen treatment experienced a 9% reduction in total thigh muscle cross-sectional area [15, 17]. To date, however, there is a complete lack of data on changes in lower-limb muscle strength during gender-affirming medical interventions. Given the established importance of muscle mass and lower-limb strength in numerous sports [18, 19], further assessment of performance indicators, as well as comprehensive muscle size and quality measures, is needed in order to provide data for sport governing bodies to stipulate informed regulations based on robust scientific evidence.

Accordingly, in the current study, we comprehensively examined the effects of gender-affirming treatment, which included inhibition of endogenous sex hormones and subsequent replacement with cross-sex hormones, on muscle function, size and composition in both TW and TM.

## Methods

### General design

This study was part of a single-center observational cohort study [20]. Examinations were conducted at 4 time points (Fig. 1): baseline (T00), 4 weeks after initiated gonadal suppression of endogenous hormones but before hormone replacement (T0), 3 months after hormone replacement therapy (T3), and 11 months after hormone-replacement therapy (T12). Each time point was divided into two separate examination days. On the first day, the participant came to the laboratory in the morning after an overnight fast. After at least 5 min of rest, blood samples were collected for standard blood biochemistry including sex hormones. On the second day, muscle strength was evaluated using isokinetic dynamometry. During the T00 and T12 visit, the participants also underwent a CT scan of the lower-limbs followed by a whole-body MRI scan. Levels of physical activity (min/week and type of activity) were assessed through questionnaires.

**Figure 1.**
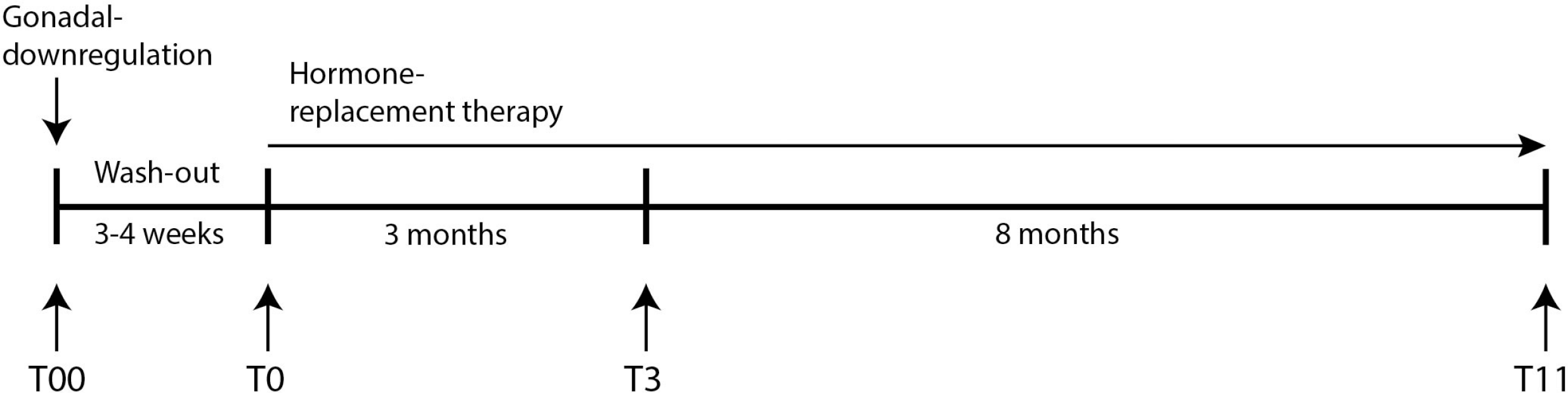
Overview of the medical intervention and examination time-points.

### Participants

The study population consisted of 12 TM (age 25±5 yrs, height 168±5 cm, body mass 66±19 kg) and 11 TW (age 27±4 yrs, height 180±5cm, body mass 70±10 kg) that were referred to Andrology Sexual Medicine and Transgender Medicine (ANOVA), at the Karolinska University Hospital, Stockholm, Sweden for evaluation of gender dysphoria. At the time of inclusion, all participants had been accepted to start gender-affirming medical interventions. Further eligibility criteria can be found in our previous publication [20] and at clinicaltrials.gov (Identifier NCT02518009). The participants were informed that their decision to participate or not, or the withdrawal of consent to participate, would not in any way change their treatment. Oral and written informed consent was obtained from all participants. The regional ethical review board in Stockholm (now called the Swedish Ethical Review Authority), approved the study (2014/409-31/4). In one of the data graphs, data from a cohort of recreationally active cis-men (n=17, age 27±5 yrs, height 181±9 cm, body mass 87±19 kg) and cis-women (n=14, age 26±4 yrs, height 163±4 cm, body mass 67±10 kg) are shown for reference. These cisgender individuals were previously assessed in our laboratory using the same measurements techniques [21].

### Medical intervention

The reversal of the endocrine environment was initiated with injection of a Gonadotropin releasing hormone (GnRH) antagonist (Degarelix 240 mg sc). This strategy is known to result in an immediate reduction in gonadotropin secretion and to bring sex hormone levels (estradiol and testosterone) to castrate levels within 24 h [22]. The subsequent cross-hormone treatment was started after post-castration assessments were made. TM were treated with testosterone injections (testosterone undecanoate 1000 mg i.m.) with the two-first injections given with a 6-week interval and thereafter one injection every tenth week, with dose adjustments in order to maintain androgen levels within the normal adult male reference range. Further gonadotropin suppression was maintained with a GnRH-analogue administered i.m. every third month. In TW, estradiol was administered transdermally (gel or patches) p.o., or in a few cases i.m.(estrogenpolyphosphate). The estradiol doses given were either 1-2 mg as gel applied daily, 100-200 μg delivered every 24 h with a patch, 4-8 mg orally, or 80 mg i.m. every 2-4 week.

### Whole-body MRI

Body composition was determined by MR imaging of the whole body at baseline and after 11 months of cross-sex hormone treatment. Each subject underwent a whole-body MR scan in the supine position (arms by their sides) using 2-point Dixon fat/water separated sequences on a 1.5 Tesla MR platform (Siemens Aera, Siemens Healthcare, Erlangen, Germany) with the following settings: slice thickness between 3.5 and 4.5 mm, repetition time 6.69, echo times 2.23 and 4.77 ms, number of averages 1, resolution 0.448 pixels per mm. The total acquisition time was <10 min. After image processing, automated image analysis was performed using segmentation software provided by AMRA™ (AMRA Medical AB, Linköping, Sweden). This analysis allowed for automated segmentation and quantification of thigh skeletal muscle volumes [23, 24], as well as total adipose tissue (TAT) and total lean tissue (TLT) volumes measured from the most distal point of the thigh muscle up to vertebrae T9, excluding arms [25]. The quantification was followed by visual quality review by a trained operator [26].

### CT-scan

The cross-sectional area and radiological density of the thigh muscles were assessed by bilateral CT scans covering the lower-limbs at baseline and after 11 months of cross-sex hormone treatment. To minimize the effect of fluid shifts, the participants rested in the supine position for 30 min before the scan [27]. Preliminary scout images were obtained to ensure accurate positioning. All scans were obtained using a second-generation 64-slice dual source CT system (SOMATOM Definition Flash, Siemens Healthcare, Forchheim, Germany) operating at 120 kV and a fixed flux of 100 mA. Areas of interest was manually defined on 5 mm thick axial slices and measured using manual planimetry with associated imaging software (Image J, National Institutes of Health, Bethesda, MD). The image selected for analysis was from the right leg, 50 mm proximal to the image where rectus femoris disappeared.

### Muscle strength assessment

Isokinetic and isometric peak torque were determined for the knee extensors and flexors using isokinetic dynamometry (Biodex System 4 Pro, Medical Systems, Shirley, NY, US) at all four time points. The chest, hip and thigh were stabilized using straps, and the ankle was strapped to the lever arm, which was aligned with the axis of rotation of the knee joint. Maximal isometric (0°/s) and isokinetic torque (60°/s and 90°/s) were measured for both the right and left leg. The participant performed 3 all-out repetitions for each leg, alternating between knee flexion and extension, at each angular velocity. A 30 s rest period was employed between the trials. For the isometric test (performed at the fixed knee angle of 120°), the participants were instructed to apply as much force as possible for 5 s. Three trials were administered for each leg interspersed with 30 s recovery.

### Blood biochemistry

The participants rested for at least 5 min prior to the blood collection. All blood samples were taken by conventional clinical procedures (sitting position, no tourniquet around the arm) from the antecubital vein. All samples were analyzed at the Karolinska University Laboratory using established procedures. The Hemoglobin measurement was based on photometry measurement of oxyhemoglobin at two wavelengths. Levels of testosterone and estradiol were measured through liquid chromatography coupled with mass-spectrometry.

### Data analysis

Data were analyzed using a 2-way repeated measures ANOVA with one group factor (TM vs. TW). The changes over time were then compared in graphs displaying the mean change with 95% confidence intervals (CI). Similarly, TW and TM were compared at the 12-month time point using a forest plot displaying the mean group difference and 95% CI. Lastly, Pearson’s R-value was computed in a linear regression analysis displaying the relationship between muscle volume and strength. The significant alpha level was set at 5%. All statistical analyses were performed using Prism 7 for Mac OS X (GraphPad Software Inc, San Diego, CA, US). Data are presented as means ± standard deviation (SD) unless otherwise stated. While we performed all lower-limb muscle volume and strength assessments for both the right and the left leg, the results confirmed that the two limbs responded very similarly and could be seen as replicates of each other. Thus, to reduce the overall volume of data presented, we report only the results from the right limb.

## Results

### Participant characteristics and blood biochemistry at baseline and 12-month follow-up

In the TM, body mass was 66±19 kg at T00 and 66±10 kg at T12 (P>0.05). In the TW, body mass was 70±10 at T00 and 73±10 at T12 (P=0.10). Total adipose tissue (TAT) volume increased from 12.7±6.8 to 16.8±6.0 L in TW (P=0.014). In contrast, TAT decreased in TM from 20.1±11.2 to 16.9±9.4 L (P=0.043). Total lean tissue (TLT) volume increased in TM from 19.1±3.1 to 21.6±3.0 L (P P<0.0001). In TW, TLT did not change significantly (P=0.15). Values were 25.0±2.4 L at T00 and 24.5±2.3 at T12. The reported levels of physical activity did not change during the intervention (P>0.05 in both groups). The reported activity consisted mainly of low- to moderate intensity aerobic activities such as brisk walking. The TM reported 216±67 min/week of physical activity at T00 and 246±128 min/week at T12. The corresponding values for the TW were 236±206 and 230±163 min/week. The blood concentrations of hemoglobin, testosterone and estradiol during the course of the study are represented in figure 2.

**Figure 2.**
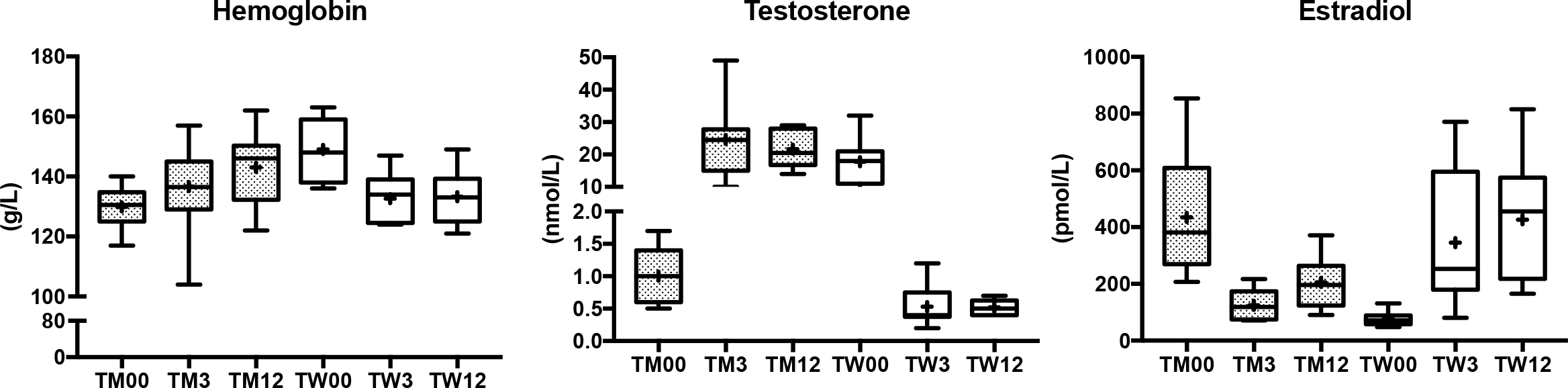
Blood biochemistry data presented as box and whisker plots. The ends of the box represent the upper and lower quartiles, while the vertical line extend to the highest and lowest observations. The median is marked by the horizontal line inside the box, and the mean value is indicated by the + symbol. TM=Transmen, TW=Transwomen. The numbers correspond to the examination time-points.

### Lower-limb muscle volume, CSA and radiological attenuation

There were group × time interactions (all P<0.0001) for all muscle volume measurements (Fig. 3). The volume of the anterior thigh (15%), posterior thigh (16%) and total thigh (15%) increased (P<0.0001) in TM. In contrast, these muscle volumes decreased (P<0.02) during the intervention in TW: anterior thigh (−5%), posterior thigh (−4%) and total thigh (−5%). There were group × time interactions (all P< 0.0001) for both quadriceps CSA and radiological density (Fig. 3). In TM, CSA increased 15%, while the radiological density increased 6%. In TW, the corresponding parameters decreased by −4% for CSA, and −2% (P>0.05) for radiological density.

**Figure 3.**
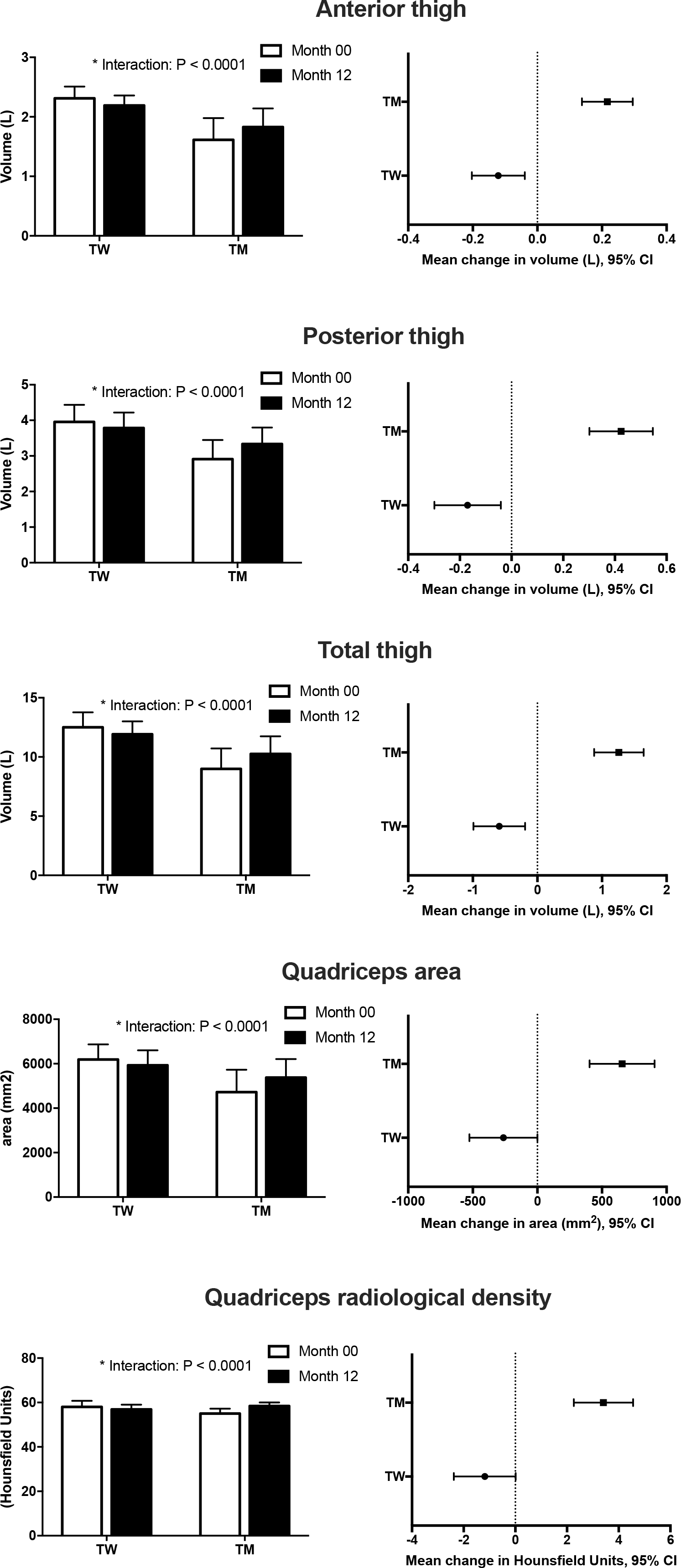
Muscle volume, determined by MRI, of the different compartments (anterior/posterior) of the right thigh, and total thigh volume (left and right together) before and after the intervention. The two bottom figures display quadriceps cross-sectional area and radiological density determined from a CT-scan. The panels to the right display the mean change over the intervention, with 95% confidence intervals. TM=Transmen, TW=Transwomen.

### Lower-limb muscle strength

Strength data and key statistics are displayed in figure 4. Isometric torque increased over the intervention for both knee extension (10%) and flexion (26%) in TM. In TW, isometric strength levels were maintained over the intervention for both extension and flexion. Isokinetic strength at 60°/s increased similarly across both TM and TW (no interactions), for both knee extension and flexion. Strength at 90°/s knee extension showed a tendency for an interaction effect (P=0.060) since strength increased in TM but was maintained in TW. For knee flexion, there was a main effect of time (P<0.0001) since both groups increased peak torque. Since strength and muscle mass increased in parallel in the TM, the specific strength (torque/quadriceps area) remained unchanged. In contrast, the specific strength increased in TW (Fig. 4).

**Figure 4.**
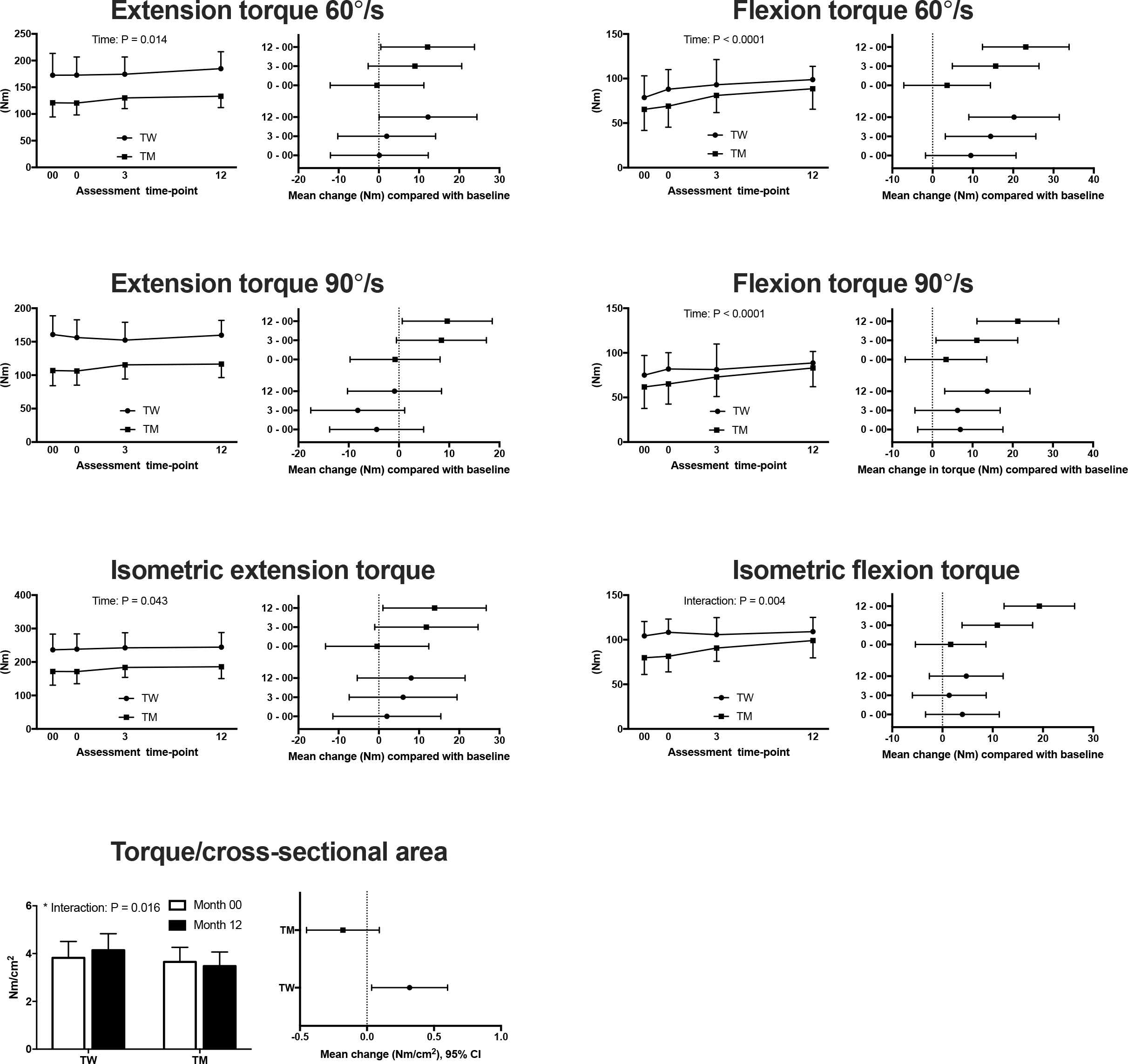
Strength data across the different velocities tested. Each strength variable is displayed with a figure showing the actual data over time (left panel of the paired figures), and the mean change across the different time-points compared (right panel of the paired figures, displayed with 95% confidence intervals of the change score in comparison with baseline T00). The two bottom figures show the isometric torque produced in relation to the quadriceps cross-sectional area (specific torque). TM=Transmen, TW=Transwomen.

### 12-month group comparison

At the 12-month visit, all lower-limb muscle volumes were still greater in TW than TM (Fig. 5). For muscle strength, all knee extension measurements favored the TW, while strength levels did not differ significantly for knee flexion (Fig. 5). While there were strong correlations (R>0.75, P<0.009) between muscle size and strength at baseline, the correlation between the change in size and change in strength was more moderate in TW (R=0.65, P=0.03), and absent in TM (R=0.32, P=0.31). Figure 6 summarizes the absolute levels of muscle strength and volume in relation to the intervention, and in comparison with the cisgender control groups.

**Figure 5.**
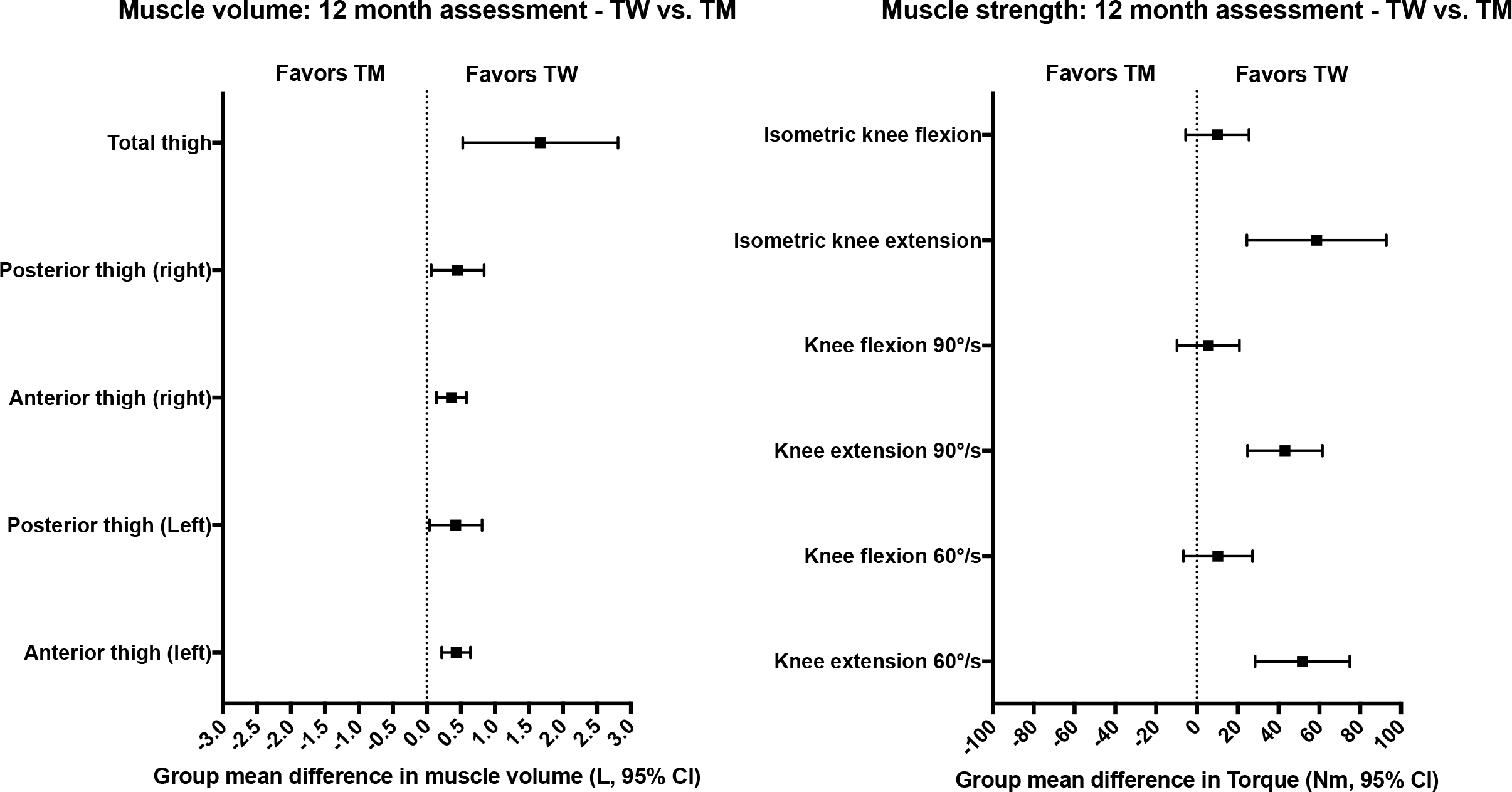
Comparison of the group mean difference (TM vs. TW) of key dependent variables at the 12-month assessment. Muscle volume measurements are displayed in the left panel, and muscle strength in the right panel. TM=Transmen, TW=Transwomen.

**Figure 6.**
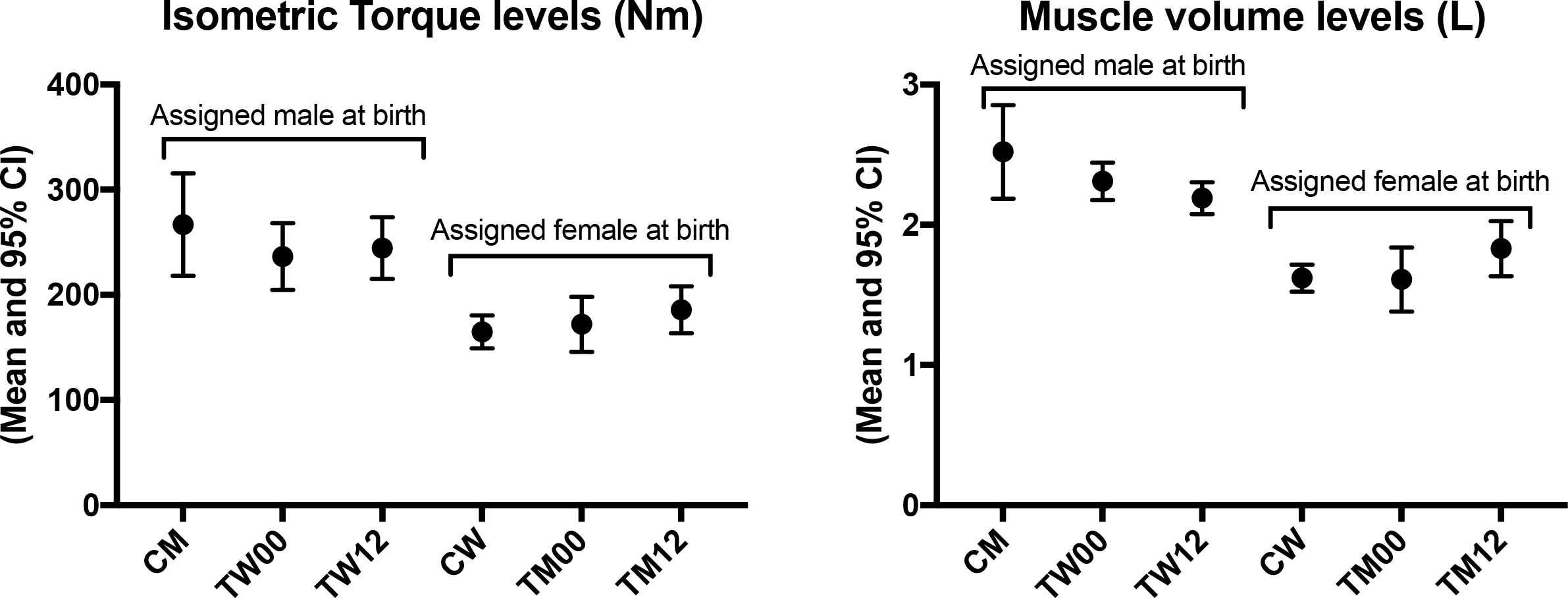
Absolute levels (with 95% confidence intervals) of isometric knee extension strength and anterior thigh muscle volume before and after the intervention, and in comparison to cis-men (n=17, CM) and cis-women (n=14, CW). TM=Transmen, TW=Transwomen.

## Discussion

The current study was conducted to comprehensively assess muscle function, size and composition in both transwomen (TW) and transmen (TM) undergoing one year of gender-affirming cross-sex hormone treatment. Our results show that thigh muscle volume increased in the TM (15%), which was paralleled by increased quadriceps cross-sectional area and radiological density. In TW, the change in muscle volume was considerably smaller, where the total loss of thigh muscle volume was 5%, which was paralleled by decreased quadriceps cross-sectional area but not radiological density. The TM increased strength over the assessment period, while the TW generally maintained or even slightly increased in strength. Despite the robust changes in lower-limb muscle mass and strength in TM, the TW still had an absolute advantage following 12 months of gender-affirming hormone treatment.

### Changes in lower-limb muscle volume, CSA and composition with 12 months of treatment

The increase in muscle volume and quadriceps CSA in the TM was expected given the well-known effect of testosterone administration on muscle mass [12, 28]. Moreover, the results and magnitudes of the gains in muscle mass were in line with previous data on TM undergoing cross-sex hormone treatment [15, 16]. A more novel finding was that the contractile density of the quadriceps also increased, as evident by the measure of radiological attenuation, a valid proxy of fat infiltration in skeletal muscle [29, 30]. This suggests that the skeletal muscles of the TM not only increase in size but also in contractile density. While this theoretically could amplify the specific tension (force produced per area) of the muscle [19], our data indicated that this did not occur in the current cohort of TM.

In contrast to the TM, the TW experienced reductions in muscle mass over the intervention. However, it is worth noting that the reduction in muscle mass in TW was smaller than the corresponding increase in TM, both in terms of relative and absolute changes. At the 12-month follow-up, the TW still had larger muscle volumes and quadriceps area than the TM. This is somewhat in contrast to two earlier reports where total thigh muscle area was similar between the TM and TW at the 12-month follow-up [15, 17]. This seems to have been driven mainly by the fact that the relative changes in muscle area in those studies were larger both in TM and TW than in the current study. Furthermore, our findings suggest that the estradiol treatment itself had a protective effect against lean mass loss, which is supported by research conducted on estrogen replacement therapy in other targeted populations [31–33] and in mice after gonadectomy [34]. The findings reported herein add important new data that increase our understanding of the effects of cross-sex hormone treatment on muscle mass and composition in transgender individuals. Overall, our results indicate that 11 months of cross-sex hormone treatment does not neutralize the pre-existing differences in muscle size between TM and TW.

### Changes in lower-limb muscle strength with 12 months of treatment

The TM increased strength over the assessment period, while the TW generally maintained or even slightly increased in strength. Given that there was no structured training performed during the intervention, the improvements in some of the strength parameters in the TW most likely arose from learning effects from repeating the test at 4 occasions, or alternatively, from a slight anabolic effect of the estrogen-based treatment [33]. In contrast, since the potential learning effect arguably would be very similar between the TM and TW, the more substantial strength increase in TM is most likely attributed to the testosterone administration [28]. At the 12-month follow-up, strength levels were still greater in the TW. This was, however, only true for knee extension and not knee flexion. Based on the data, the most obvious explanation for this was the relatively smaller differences between groups already at baseline for knee flexion strength, and the more substantial increase in knee flexion compared to knee extension strength in the group of TM.

### Methodological considerations

We acknowledge that this study was conducted with untrained individuals and not transgender athletes. Thus, while this gave us the important opportunity to study the effect of the cross hormone treatment alone, and as such the study adds important data to the field, it is still uncertain how the findings would translate to transgender athletes undergoing advanced training regimens during the gender-affirming intervention. It is also important to recognize that we only assessed proxies for athletic performance, such as muscle mass and strength. Future studies are needed to examine a more comprehensive battery of performance proxies in transgender athletes. Given the marked changes in hemoglobin concentration in the current study, it is possible that gender-affirming treatment also has effects on endurance performance and aerobic capacity. Furthermore, since the TM and to some extent also the TW demonstrated progressive changes in muscle strength, and strikingly some of the TW individuals did not lose any muscle mass at all, follow-ups longer than 12 months are needed to better characterize the long-term consequences and individual responsiveness to gender affirming interventions.

### Conclusions and implications

Cross-sex hormone treatment markedly affects muscle function, size and composition in transgender individuals. Despite the robust changes in lower-limb muscle mass and strength in TM, the TW still had an absolute advantage at the 12-month follow-up. The question of when it is fair to permit a Transwoman to compete in sport in line with her experienced gender identity is challenging and very little data have been provided to add clarity on the potential performance benefits for TW arising from the lifelong experience of being a biological male. Our results indicate that after 12 months of hormonal therapy, a transwoman will still likely have performance benefits over a cis-woman. These findings add new knowledge that could be relevant for sport-governing bodies when evaluating the eligibility of transwomen to compete in the women’s category of athletic competition.

## Contribution

Conception and design of the study: TG, SA, AW. Data acquisition, analysis or interpretation of data: TRL, AW, ER, MM, DPA, TBB, ODL, JF, MH, SC. Drafting the manuscript: AW, TRL. All authors revised the manuscript critically for important intellectual content and approved the final version.

## Ethics approval

The Regional Ethical Review Board in Stockholm (now called the Swedish Ethical Review Authority, No. 2014/409-31/4)

